# *Patolakaturohiniyadi Kashayam*, exerts anti-steatotic and anti-obesogenic effects via coordinated regulation of lipid metabolism, inflammation, and incretin signalling

**DOI:** 10.64898/2026.07.14.738366

**Authors:** Sania Kouser, Subrahmanya Kumar, Poornima Devkumar, Chethala N Vishnuprasad

**Author notes:** Corresponding author: Chethala N Vishnuprasad.

## Abstract

**Background:** Metabolic dysfunction is characterized by dysregulated lipid metabolism, lipotoxicity, insulin resistance, and chronic low-grade inflammation, contributing to obesity and metabolic dysfunction-associated steatotic liver disease (MASLD). Multi-target therapeutic strategies that restore lipid homeostasis are of growing interest. *Patolakaturohiniyadi Kashayam* (PKR), a classical Ayurvedic polyherbal formulation, was investigated for its potential to modulate lipid metabolism and ameliorate metabolic dysfunction.

**Methods:** An integrated approach combining network pharmacology, in vitro, lipidomics, and in vivo studies was employed. Hub gene identification and KEGG pathway enrichment were performed to elucidate molecular targets. Anti-steatotic and anti-adipogenic effects were assessed in hepatocytes and adipocytes, followed by lipidomic profiling. Efficacy was further evaluated in a high-fat high-fructose diet (HFHFD)-induced animal model.

**Results:** Network pharmacology identified key targets including TP53, AKT1, IL6, TNF, and STAT3, enriched in pathways related to lipid metabolism, inflammation, and metabolic regulation. PKR significantly reduced lipid droplet accumulation and intracellular triglyceride levels in vitro. Lipidomics revealed suppression of diacylglycerol-mediated lipotoxicity and restoration of phospholipid balance, characterized by increased lysophospholipids and phosphatidylethanolamines with normalization of phosphatidylcholine species. In vivo, PKR reduced body, liver, and adipose tissue weights, improved serum lipid profiles, and decreased AST and ALT levels. Histological analyses demonstrated reduced lipid accumulation and inflammation, along with preservation of adipose tissue architecture. PKR also improved glucose tolerance and significantly elevated plasma GLP-1 levels.

**Conclusion:** PKR exerts potent anti-steatotic and anti-obesogenic effects through coordinated regulation of lipid metabolism, inflammation, and incretin signalling, highlighting its potential as a multi-target therapeutics for metabolic dysfunction.

## 1. Introduction

Obesity has emerged as a major global public health concern, driven by rapid urbanization, sedentary lifestyles, and increased consumption of calorie-dense diets. According to the World Health Organization, more than 1.9 billion adults are overweight, of which over 650 million are obese, contributing significantly to global morbidity and mortality (1). In India, the burden of obesity has increased dramatically over the past few decades, with an estimated 135 million individuals affected and prevalence rates ranging from 11.8% to 31.3% across different regions (2). Recent epidemiological studies further indicate a rising prevalence of metabolic abnormalities even among non-obese individuals, reflecting a growing trend of metabolically unhealthy phenotypes in the Indian population (3).

Closely associated with obesity is metabolic dysfunction-associated steatotic liver disease (MASLD) (previously referred to as NAFLD, Non-alcoholic Fatty Liver Disease), which represents the hepatic manifestation of systemic metabolic imbalance. MASLD is estimated to affect approximately 25–30% of the global population, with prevalence in India reported to be as high as 30–40%, particularly among individuals with obesity, type 2 diabetes, and dyslipidemia (4,5). Importantly, obesity is a major risk factor for MASLD, and both conditions share common pathophysiological mechanisms, including dysregulated lipid metabolism, insulin resistance, and chronic low-grade inflammation. Together, they form a metabolic continuum that substantially increases the risk of cardiovascular disease and other complications, thereby contributing to a significant healthcare burden (6).

At the mechanistic level, adipose tissue plays a central role in the development of metabolic dysfunction. Excess caloric intake promotes adipocyte hypertrophy and hyperplasia through enhanced adipogenesis, resulting in increased lipid storage. However, chronic overexpansion of adipose tissue leads to adipocyte dysfunction, characterized by hypoxia, inflammation, and impaired lipid buffering capacity (7). This dysfunction results in elevated circulating free fatty acids and ectopic lipid deposition in peripheral organs such as the liver. Consequently, hepatic steatosis develops due to increased lipid influx, enhanced de novo lipogenesis, and reduced fatty acid oxidation (8). Furthermore, accumulation of lipotoxic intermediates such as diacylglycerols exacerbates insulin resistance and activates inflammatory signaling pathways, thereby perpetuating metabolic dysfunction (9).

Despite the increasing prevalence and disease burden of obesity and MASLD, current therapeutic options remain limited. Lifestyle interventions, including dietary modification and physical activity, are the cornerstone of management but are often difficult to sustain long term (10). Pharmacological therapies largely target individual metabolic parameters such as hyperglycemia or dyslipidemia and fail to address the complex, interconnected nature of metabolic dysfunction. Moreover, several drugs are associated with suboptimal efficacy and potential adverse effects, highlighting the need for safe and effective multi-target therapeutic strategies that can modulate lipid metabolism, inflammation, and metabolic homeostasis simultaneously (11).

In this context, traditional herbal formulations have gained increasing attention due to their multi-component and pleiotropic mechanisms of action. *Patolakaturohiniyadi Kashayam* (PKR) is a classical Ayurvedic polyherbal formulation traditionally used in the management of metabolic and hepatobiliary disorders. Its constituent herbs are reported to possess hepatoprotective, anti-inflammatory, and metabolic regulatory properties, suggesting potential efficacy in modulating lipid metabolism and systemic metabolic dysfunction (Table1) (12). However, its mechanistic basis and therapeutic potential remain inadequately explored in the context of lipid dysregulation and obesity-associated metabolic imbalance. Therefore, the present study aimed to investigate the anti-steatotic and anti-obesogenic effects of PKR using an integrated approach combining network pharmacology, in vitro assays, lipidomics, and in vivo validation. The study further sought to elucidate the molecular mechanisms underlying PKR action, with a particular focus on lipid metabolism, lipotoxicity, and metabolic regulation, to establish its potential as a multi-target therapeutic strategy for metabolic dysfunction.

## 2. Materials and methods

### 2.1. Chemicals and reagents

Dulbecco’s modified Eagle’s medium; fetal bovine serum (FBS); and penicillin/streptomycin (PS) (GIBCO, Grand Island, New York); Non-esterified fatty acid free bovine serum albumin (BSA) (Genei, Bangalore, India); triglyceride kit (BeneSphera, Avantor, USA); ascorbic Acid; Folin-Ciocalteu reagent; gallic acid; sodium palmitate (PA); Oil-Red-O (ORO) stain; IBMX; dexamethasone; insulin (Sigma-Aldrich, St. Louis, MO, USA); SPLASH^®^ LIPIDOMIX^®^ Mass Spec Standard (Avanti polar lipids, Alabaster, AL, USA, 330707); Pioglitazone hydrochloride tablet (USV Private Limited, Mumbai, India, PIOZ 15); GLP-1 kit (RayBiotech Life Inc., Peachtree Corners, GA, USA, EIA-GLP1); Insulin Elisa kit (Krishgen biosystems, Mumbai, India, KLR0707); AST kit (Elabscience, Texas, USA, E-BC-K236-M); ALT kit (Elabscience, Texas, USA, E-BC-K235-M).

### 2.2. Network Pharmacology

A systematic network pharmacology approach was employed to elucidate the multi-component, multi-target mechanisms underlying the therapeutic potential of the selected polyherbal formulation. Initially, phytochemical constituents of the six medicinal plants were retrieved from publicly available databases, including IMPPAT and Dr. Duke’s Phytochemical and Ethnobotanical Databases. All identified compounds were curated, and duplicates were removed to obtain a final dataset of 41 unique phytochemicals, which were considered for subsequent analysis. To predict potential molecular targets, each phytochemical was queried against multiple chemoinformatics databases, including ChEMBL, STITCH, and BindingDB. The retrieved targets were compiled, standardized to gene symbols, and de-duplicated, resulting in a total of 333 unique target genes associated with the selected phytochemicals. A compound–target interaction network was constructed using Cytoscape (version 3.10.4), where nodes represented phytochemicals and target genes, and edges denoted their predicted interactions. This network enabled visualization of the complex relationships between multiple bioactive compounds and their corresponding targets. To further investigate the functional interactions among the identified targets, a protein–protein interaction (PPI) network was generated using the STRING database. The list of 333 genes was uploaded with the organism restricted to *Homo sapiens*, and a high-confidence interaction score (>= 0.7) threshold was applied to ensure reliability of the predicted associations. The resulting PPI network was imported back into Cytoscape for visualization and topological analysis. Identification of key regulatory genes within the network was performed using the CytoHubba plugin in Cytoscape. Multiple topological parameters, including degree centrality, maximal clique centrality (MCC), and betweenness centrality, were employed to rank nodes based on their importance within the network. Genes consistently ranked highly across these parameters were considered as hub genes and were selected for downstream analysis. Subsequently, functional enrichment analysis of the selected hub genes was conducted using the ClueGO plugin within Cytoscape. Kyoto Encyclopedia of Genes and Genomes (KEGG) pathway enrichment analysis was performed to identify significantly associated biological pathways. Statistical significance was determined using a two-sided hypergeometric test, followed by Benjamini–Hochberg correction for multiple testing. Pathways with adjusted *p* < 0.05 were considered significant.

### 2.3. PKR quantification

PKR was procured from an authorized *Ayurveda* drug manufacturer (Kottakkal Arya Vaidya Sala, Kerala) with batch numbers 530236 and 529747. The total phenolic content was estimated following Folin - Ciocalteu method, using gallic acid as standard, and the experimental concentrations were expressed as microgram of gallic acid equivalent per milliliter (μg GAE/mL) (13, 14).

### 2.4. Mammalian cell maintenance and modelling of liver steatosis and adipogenesis

HepG2 (human hepatoma cell line, Passage 25-35) and 3T3-L1 (mouse fibroblasts cell line, Passage 10-17) were procured from National Centre for Cell Sciences (NCCS), Pune, India, and were maintained in DMEM containing 10% FBS and 1% PS in a humidified atmosphere containing 5% CO_2_ at 37 °C. To establish a palmitic acid induced steatosis model in HepG2 cells, the cells grown in multi-well plates for 48 h were induced with 1 mM palmitic acid (PA) in serum-free DMEM containing 5% BSA for 24 h. After induction, the cells were treated with 30 µg GAE/mL PKR prepared in the same BSA containing medium for 24 h to assess the anti-steatotic effect of the formulation. For inducing adipogenesis, two days post-confluent (day-0) 3T3-L1 fibroblast cells were grown in differentiation induction medium (MDI – growth media containing 500 μM IBMX, 250 nM dexamethasone and 50 nM insulin) for three days, followed by insulin media (growth media + 50 nM insulin) for 2 days and subsequently the cells were maintained in fresh culture media till they attain complete adipocyte morphology. To assess the anti-adipogenic effect of PKR, the same induction protocol was followed with an addition of 4 µg GAE/mL PKR along with MDI treatment.

### 2.5. Oil-Red-O staining of intracellular lipid accumulation and triglyceride accumulation in HepG2 and 3T3-L1 cells

Briefly, HepG2 (for steatosis) and 3T3-L1 cells (for adipogenesis) were washed twice with PBS, and fixed with 4% formaldehyde for 30 min. After washing, cells were stained for 20 min in freshly diluted ORO solution (0.5% ORO in isopropanol diluted to 3:2 with H_2_O, and filtered) at 37°C. The unbound stain was removed by thoroughly washing the cells with double distilled water and images of cells stained with Oil Red were captured using phase contrast microscope (Olympus-IX-Olympus America Inc, USA). For quantitation, ORO stain was eluted with 200 µL isopropanol and the optical density was measured at 450 nm using a microplate reader.

To analyze the cellular triglyceride content, both HepG2 and 3T3-L1 cells, after drug treatment, were washed with PBS and scraped into 250 μL of 0.1% PBST and pulse sonicated at 45% amplitude for 45 s. The lysates were assayed using triglyceride assay kit and normalized to total protein. The results were expressed as µg of triglyceride per mg of cellular protein.

### 2.6. Global lipidomic analysis

Treated HepG2 cells were washed thrice with ice cold PBS and lipids were extracted following Bligh and Dyer method with slight modification (15). To the cell pellet 10 μl of internal standard was added followed by sequential addition of 200 µl each of ice-cold methanol, chloroform and 0.8% KCl. The samples were pulse sonicated (1 min), incubated (37 °C, 10 min) and centrifuged at 10,000 rpm for 10 min (4 °C). The lower organic phase was collected and dried using speed vac, and reconstituted in 100 µl of methanol. 5 µl of the sample was injected into the Orbitrap mass spectrometer for lipidomic analysis. Data were aligned by MS-DIAL software and statistical analysis was done using Metaboanalyst 6.0. The obtained abundance data was initially filtered using IQR threshold of 25% with median intensity applied as abundance filter. Data was normalized using log_10_ data transformation followed by auto-scaling. A p-value of < 0.05 was considered statistically significant for all analysis. Pathway enrichment was carried out using the BioPAN database.

### 2.7. *In Vivo* studies

Male Sprague-Dawley rats (SD rats) weighing 180-200 g were procured and maintained in the animal house facility of Acharya & BM Reddy college of Pharmacy, Bengaluru, India; under controlled environmental conditions (temperature 22±2 °C; relative humidity 50±5% and 12 h light and dark cycle). Experiments were conducted following approved protocols and guidelines outlined by the Committee for Control and Supervision of Experiments on Animals (CCSEA), Government of India (IAEC/ABMRCP/2023-2024/4) and the ARRIVE guidelines. An acute toxicity study was performed following the limit test standard of the Organization for Economic Cooperation and Development (OECD) No. 425 Guideline and found that the formulation is well tolerated with no fatality or signs of abnormal behavior (16).

### 2.8. Induction of MASLD in experimental rats

After acclimatization, the initial weight and blood glucose levels of animals were recorded and were randomly divided into two nutritional groups: a) Control group that received standard chow diet (3060 kcal/kg) and b) HFHFD group that received high-fat-high-fructose diet (5200 kcal/kg) (30% palm oil, 20% fructose, 2% cholesterol and 0.5% cholic acid). Commercial HFHF diet was procured from VRK nutritional solutions, Pune, India. (Supplementary Data 1). Both the groups received normal drinking water *ad libitum*. Daily food intake and weekly body weight were recorded throughout the study. After 90 days, the HFHFD group were randomly assigned to experimental groups (Table 4) receiving oral administration of either the formulation or placebo, once in a day for 30 days, while maintaining the HFHFD regimen. The doses were calculated according to the rat equivalent volume of adult human dose, derived by multiplying the adult human dose by 0.018 per 200 g of body weight of rat (17). The adult human dose of formulations was considered to be 30 mL as high dose and 15 mL as low dose as per *Ayurveda* clinical practice.

### 2.9. Oral Glucose Tolerance Test (OGTT)

At the end of drug intervention, the rats were fasted overnight, and the basal blood glucose level was recorded. The different experimental groups were administered with 2 g/kg of glucose orally and the blood glucose was measured at 15, 30, 60 and 120 min by making an incision at the caudal vein at the tip of the tail, using a commercial glucometer (Accucheck Active). The results were expressed as the integrated area under the curve for glucose (AUC glucose).

### 2.10. Biochemical parameter assessment and histopathological analysis

At the end of the experiment, blood was collected from the retro-orbital sinus into two tubes, and the rats were euthanized under over-dose of anesthesia for tissue collection. The blood was centrifuged at 3500 rpm for collecting plasma and serum for various biochemical analysis following the manufacturer’s protocol. The liver, adipose and pancreas tissues from each group were immediately removed and fixed in 10% buffer saturated formalin and were subjected to histological analysis (Dr Vamshi’s Biological Sciences and Research Center, Rajajinagar, Bangalore).

### 2.11. Statistical analysis

The results were expressed as the mean□±□standard error of the mean (SEM). Statistical significance was assessed using the analysis of variance (ANOVA) with post hoc Bonferroni correction for multiple comparisons. All statistical analyses were conducted using GraphPad Prism software (version 10.6.1). Values of *p□*<□0.05 (*) or□<□0.005 (**) or <□0.001 (***) were considered statistically significant.

## 3. Results

### 3.1. Network Pharmacology Analysis Reveals Key Hub Genes and Pathways Associated with Metabolic and Inflammatory Regulation

The constructed interaction network demonstrated a highly interconnected architecture, reflecting the multi-target nature of the formulation. Several target genes exhibited high connectivity, indicating their central role in mediating the biological effects of the phytochemical constituents. Topological analysis of the protein–protein interaction network identified a subset of highly influential hub genes based on integrated centrality parameters (Table 2). Among these, TP53, EGFR, TNF, IL6, STAT3, AKT1, and IL1B emerged as core regulators within the network. These genes are primarily associated with inflammatory signalling, immune modulation, and cellular survival processes, underscoring their importance in metabolic dysfunction. In addition to these central regulators, a distinct cluster of genes involved in xenobiotic metabolism and hepatic function was observed, including CYP3A4, CYP2C9, UGT1A1, and UGT1A6. The enrichment of these genes suggests a potential role in regulating hepatic detoxification and maintaining metabolic homeostasis. Functional enrichment analysis further demonstrated that the hub genes were significantly associated with pathways related to metabolic dysfunction and inflammation (Table 3). Prominent pathways included lipid and atherosclerosis, IL-17 signalling, TNF signalling, and AGE–RAGE signalling in diabetic complications. Importantly, enrichment of the adipokine signalling pathway highlights the role of these targets in regulating adipose tissue function, insulin sensitivity, and energy homeostasis. Notably, the enrichment of the non-alcoholic fatty liver disease (NAFLD (MASLD)) pathway further establishes a direct link between the identified targets and hepatic lipid accumulation, inflammation, and disease progression. These findings indicate that the therapeutic effects are closely aligned with key pathological processes underlying metabolic liver disorders. Additional enrichment in pathways associated with immune activation, including toll-like receptor signalling, further supports the involvement of innate immune responses and chronic low-grade inflammation. The presence of infection-and cancer-related pathways likely reflects shared molecular mechanisms such as cytokine signalling, oxidative stress, and cellular proliferation. Collectively, these results highlight a coordinated network in which key inflammatory mediators, adipokine-regulated metabolic signals, and hepatic detoxification enzymes converge to modulate disease-associated pathways. This integrative mechanism suggests that the formulation exerts its effects through simultaneous regulation of inflammation, adipose tissue signalling, and liver metabolism, thereby targeting multiple aspects of metabolic dysfunction.

**Table 1.** Plant composition of PKR.

| Sanskrit Name | Scientific Name | Part used |
| --- | --- | --- |
| Patola | Trichosanthes lobata | Pl. |
| Katurohini | Neopicrorhiza scrophulariiflora | Rt. |
| Chandana | Santalum album | Ht. Wd. |
| Madhusrava | Moringa oleifera | Rt. |
| Guluchi | Tinospora cordifolia | St. |
| Patha | Cyclea peltata | Rt. |

**Table 2.**
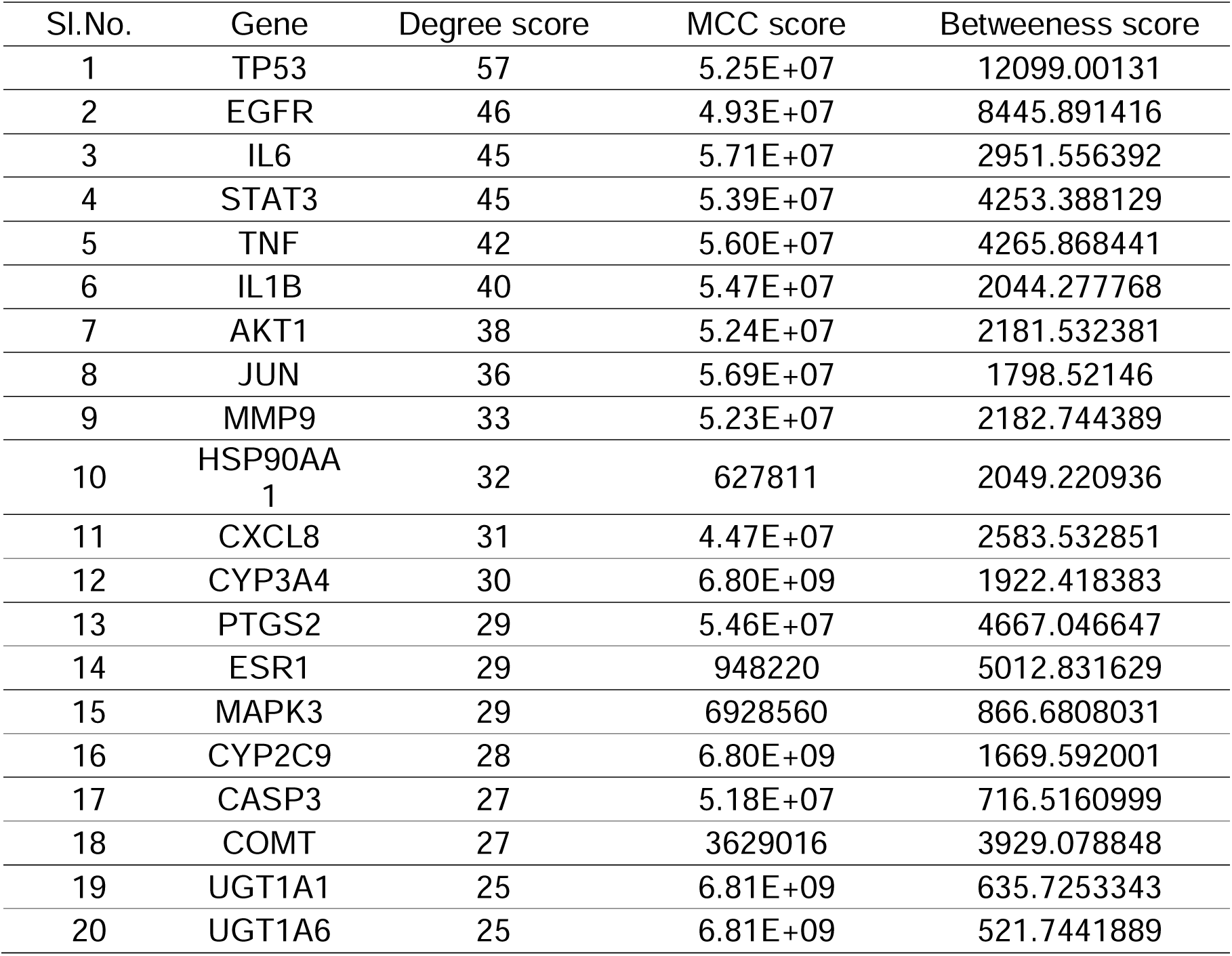
Top 20 hub genes identified from the protein–protein interaction network based on degree centrality, with corresponding MCC and betweenness centrality scores.

| Sl.No. | Gene | Degree score | MCC score | Betweenness score |
| --- | --- | --- | --- | --- |
| 1 | TP53 | 57 | 5.25E+07 | 12099.00131 |
| 2 | EGFR | 46 | 4.93E+07 | 8445.891416 |
| 3 | IL6 | 45 | 5.71E+07 | 2951.556392 |
| 4 | STAT3 | 45 | 5.39E+07 | 4253.388129 |
| 5 | TNF | 42 | 5.60E+07 | 4265.868441 |
| 6 | IL1B | 40 | 5.47E+07 | 2044.277768 |
| 7 | AKT1 | 38 | 5.24E+07 | 2181.532381 |
| 8 | JUN | 36 | 5.69E+07 | 1798.52146 |
| 9 | MMP9 | 33 | 5.23E+07 | 2182.744389 |
| 10 | HSP90AA1 | 32 | 627811 | 2049.220936 |
| 11 | CXCL8 | 31 | 4.47E+07 | 2583.532851 |
| 12 | CYP3A4 | 30 | 6.80E+09 | 1922.418383 |
| 13 | PTGS2 | 29 | 5.46E+07 | 4667.046647 |
| 14 | ESR1 | 29 | 948220 | 5012.831629 |
| 15 | MAPK3 | 29 | 6928560 | 866.6808031 |
| 16 | CYP2C9 | 28 | 6.80E+09 | 1669.592001 |
| 17 | CASP3 | 27 | 5.18E+07 | 716.5160999 |
| 18 | COMT | 27 | 3629016 | 3929.078848 |
| 19 | UGT1A1 | 25 | 6.81E+09 | 635.7253343 |
| 20 | UGT1A6 | 25 | 6.81E+09 | 521.7441889 |

**Table 3.**
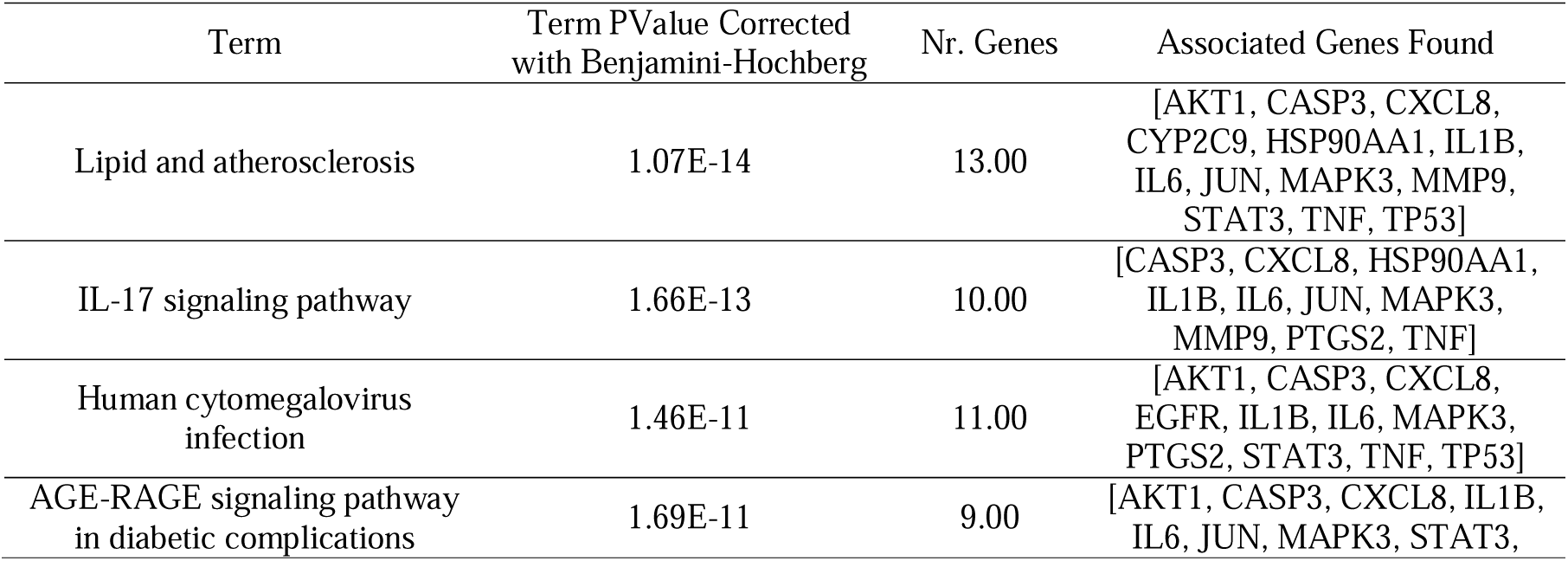
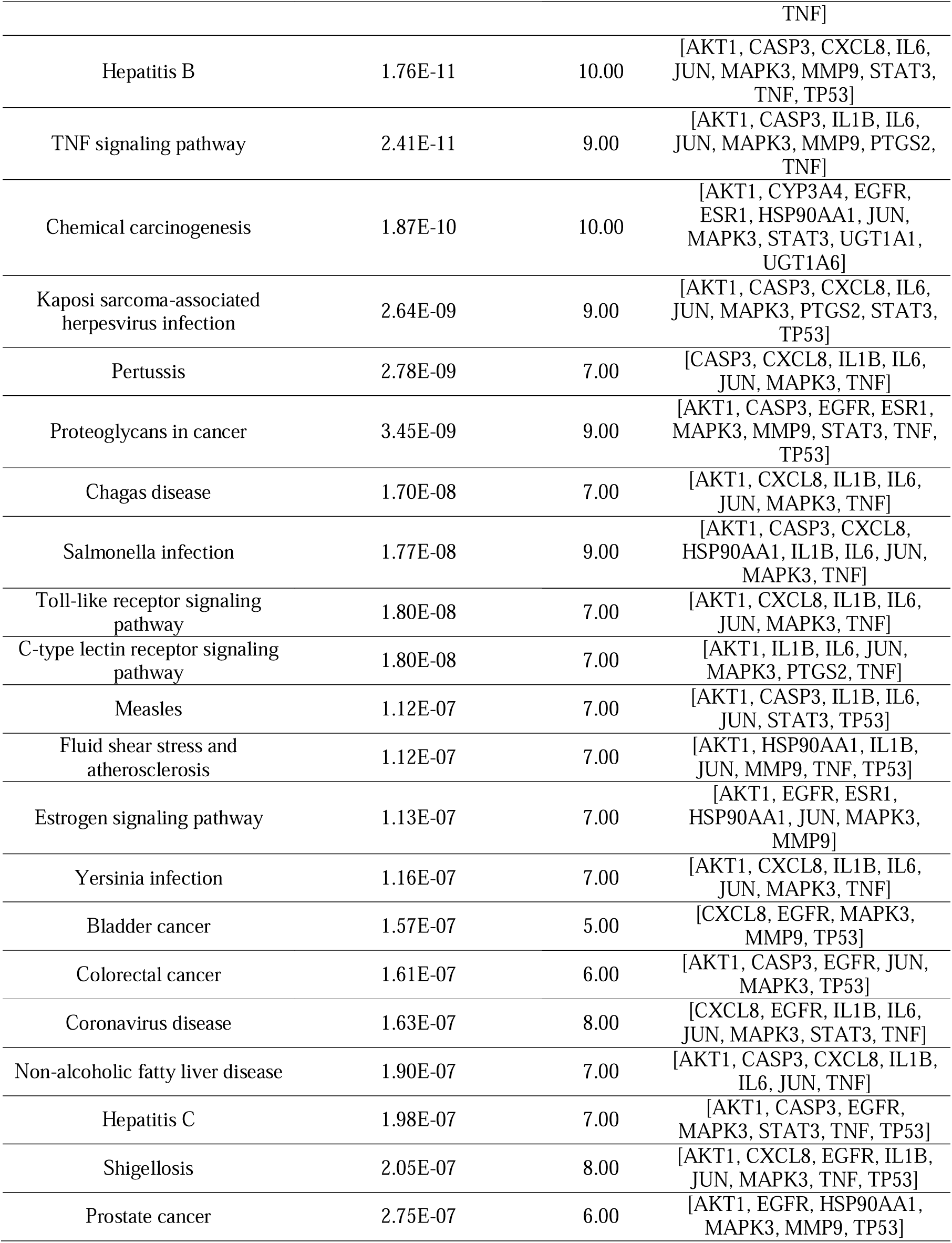
Functional enrichment of hub genes highlighting key metabolic, inflammatory, and disease-associated KEGG pathways.

**Table 4:** Details of animal grouping and treatment administered.

| Sl. No. | Group | Treatment | No. of animals |
| --- | --- | --- | --- |
| 1 | Control | Placebo | 5 |
| 2 | HFHFD | Placebo | 5 |
| 3 | PIOZ | 0.6 mg/200 g/day | 5 |
| 4 | PKRHD | 0.27 mL/200 g/day | 5 |
| 5 | PKRLD | 0.135 mL/200 g/day | 5 |

### 3.2. PKR Attenuates Lipid Accumulation in Hepatocytes and Adipocytes

Palmitic acid (PA) treatment induced significant lipid accumulation in both hepatocytes and adipocytes, as evidenced by increased intracellular lipid droplet formation and elevated triglyceride levels. In hepatocytes, PA exposure resulted in pronounced steatosis, characterized by dense and enlarged lipid droplets. Similarly, adipocytes exhibited enhanced lipid storage, reflecting increased adipogenic activity.

Treatment with PKR markedly reduced lipid accumulation in both cell types. Microscopic analysis revealed a substantial decrease in the number and size of lipid droplets following PKR treatment compared to PA-treated controls. This reduction was consistently observed across hepatocytes and adipocytes, indicating a broad-spectrum effect on lipid handling. Quantitative assessment further demonstrated a significant decline in intracellular triglyceride levels upon PKR treatment in both models. In hepatocytes, this suggests an attenuation of lipid overload and steatotic progression, while in adipocytes, it reflects suppression of lipid storage and adipogenic processes. Collectively, these findings indicate that PKR exerts a dual anti-steatotic and anti-adipogenic effect, effectively reducing lipid accumulation under lipotoxic conditions. This highlights its potential to modulate key processes involved in metabolic dysfunction, including hepatic steatosis and adipose tissue lipid storage.

### 3.3. PKR reprograms the hepatocellular lipidome by attenuating DG-driven lipotoxicity and restoring phospholipid homeostasis

Untargeted lipidomic profiling was performed to elucidate the impact of PKR on hepatocellular lipid remodelling under lipotoxic conditions. In the positive ion mode, PKR treatment resulted in a marked increase in lysophospholipid species, including nine lysophosphatidylcholines (LPCs) and two lysophosphatidylethanolamines (LPEs), with notable elevation of LPC (16:0), LPC (18:0), and LPE (20:1). Concurrently, PKR normalized key phosphatidylcholine (PC) species, including PC (36:4) and PC (38:9), indicating restoration of membrane phospholipid balance. Importantly, PKR significantly suppressed diacylglycerol (DG) species, particularly DG (16:0_16:0) and DG (16:1_16:2), which are established mediators of lipotoxicity and insulin resistance. In the negative ion mode, PKR predominantly modulated phosphatidylethanolamine (PE)-rich membranes, leading to a substantial increase in multiple PE species, including PE (16:1_18:1), PE (16:0_16:1), PE (18:0_20:3), PE (18:0_22:5), and PE (18:1_22:6). Additionally, PKR influenced selected PC species, such as PC (18:1_18:2) and PC (18:2_17:2;O), further supporting its role in phospholipid remodelling. Collectively, these findings demonstrate that PKR effectively restores the palmitic acid (PA)-altered hepatocellular lipidome by suppressing DG-driven lipotoxicity and reprogramming phospholipid composition. The shift toward increased lysophospholipids and PE enrichment, along with normalization of PC species, suggests enhanced membrane dynamics and improved cellular homeostasis. This lipidomic reconfiguration highlights a hepatoprotective mechanism of PKR, characterized by stabilization of membrane integrity and mitigation of lipotoxic stress.

### 3.4. PKR alleviates systemic metabolic burden and restores hepatic and adipose tissue integrity

Administration of PKR resulted in a pronounced improvement in systemic metabolic parameters, as evidenced by a significant reduction in body weight, liver weight, and adipose tissue mass across all treatment groups compared to the HFHFD control (Fig. 4 B–D). These findings indicate a robust anti-obesogenic effect, reflecting improved whole-body metabolic homeostasis. Consistent with these observations, PKR markedly improved the circulating lipid profile. All treatment groups exhibited significant reductions in serum triglycerides, total cholesterol, and very-low-density lipoprotein (VLDL) levels (Fig. 4 H–J), highlighting its efficacy in mitigating dyslipidemia. Notably, the high-dose PKR group demonstrated a significant elevation in high-density lipoprotein (HDL) levels (Fig. 4 K), suggesting enhanced lipid transport and cardioprotective potential. PKR treatment also conferred significant hepatoprotective effects, as reflected by the substantial reduction in serum hepatic injury markers, including aspartate aminotransferase (AST) and alanine aminotransferase (ALT) (Fig. 4 L–M). Histopathological evaluation further corroborated these biochemical findings. Liver sections from PKR-treated groups exhibited a marked attenuation of macrovesicular steatosis, lobular inflammation, hepatocyte ballooning, and fibrosis compared to the HFHFD group (Fig. 4 E–F, 400×), indicating a reversal of diet-induced hepatic injury. In parallel, adipose tissue histology revealed well-preserved adipocyte architecture in PKR-treated animals, with an absence of vascular congestion and inflammatory infiltration (Fig. 4 G). This preservation of adipose tissue integrity underscores the ability of PKR to prevent adipose dysfunction and associated inflammatory responses. Collectively, these findings demonstrate that PKR exerts comprehensive metabolic benefits by reducing adiposity, improving lipid profiles, and restoring both hepatic and adipose tissue structure, thereby mitigating systemic metabolic stress.

**Figure 1.**
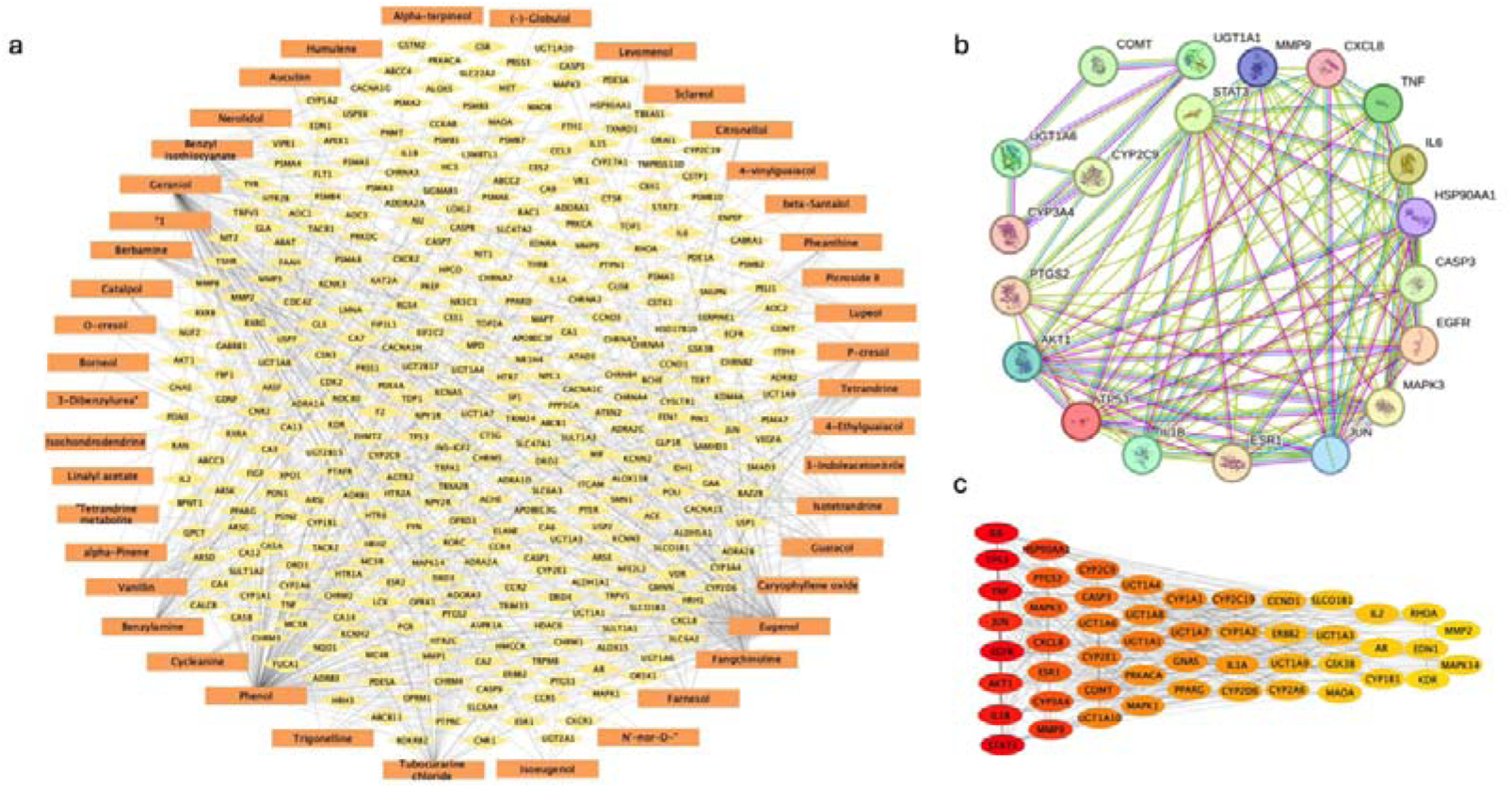
Network Pharmacology Analysis Reveals Key Hub Genes and Pathways Associated with Metabolic and Inflammatory Regulation. a) Network diagram of PKR phytoconstituents (orange) with target genes (yellow), b) PPI network of hub genes, c) Network of Hub genes based on degree of betweenness.

**Figure 2.**
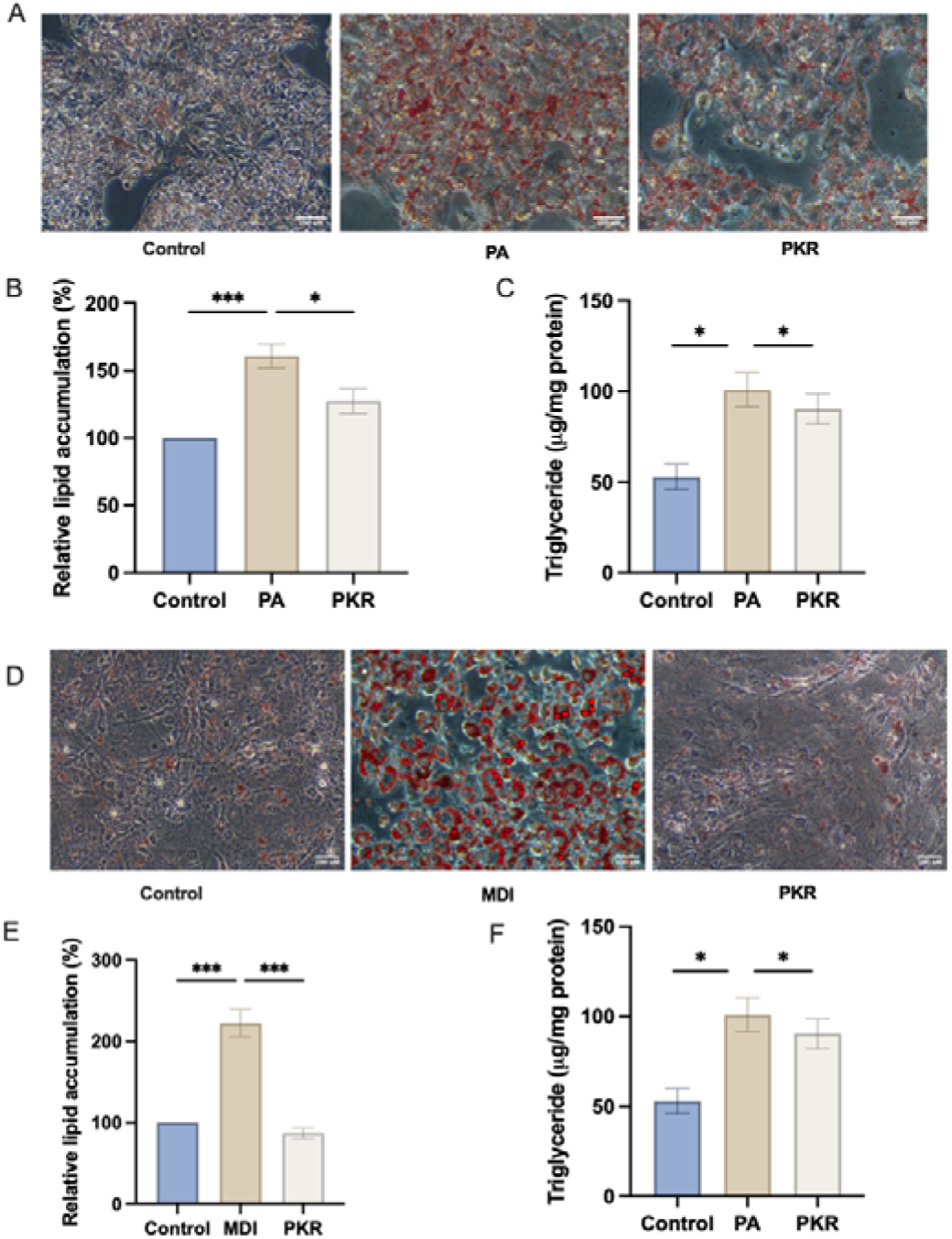
PKR attenuates lipid accumulation in PA-induced in hepatocytes and MDI-induced adipocytes. (A-B) Oil Red O staining (20x) and quantification of intracellular lipid content in hepatocytes; (C) Triglyceride (TG) assay in hepatocytes; (D-E) Oil Red O staining (20x) and quantification of intracellular lipid content in adipocytes; (F) Triglyceride (TG) assay in adipocytes. Data are presented as mean□±□SEM; *p□<□0.05, **p□<□0.005, ***p□<□0.001.

**Figure 3.**
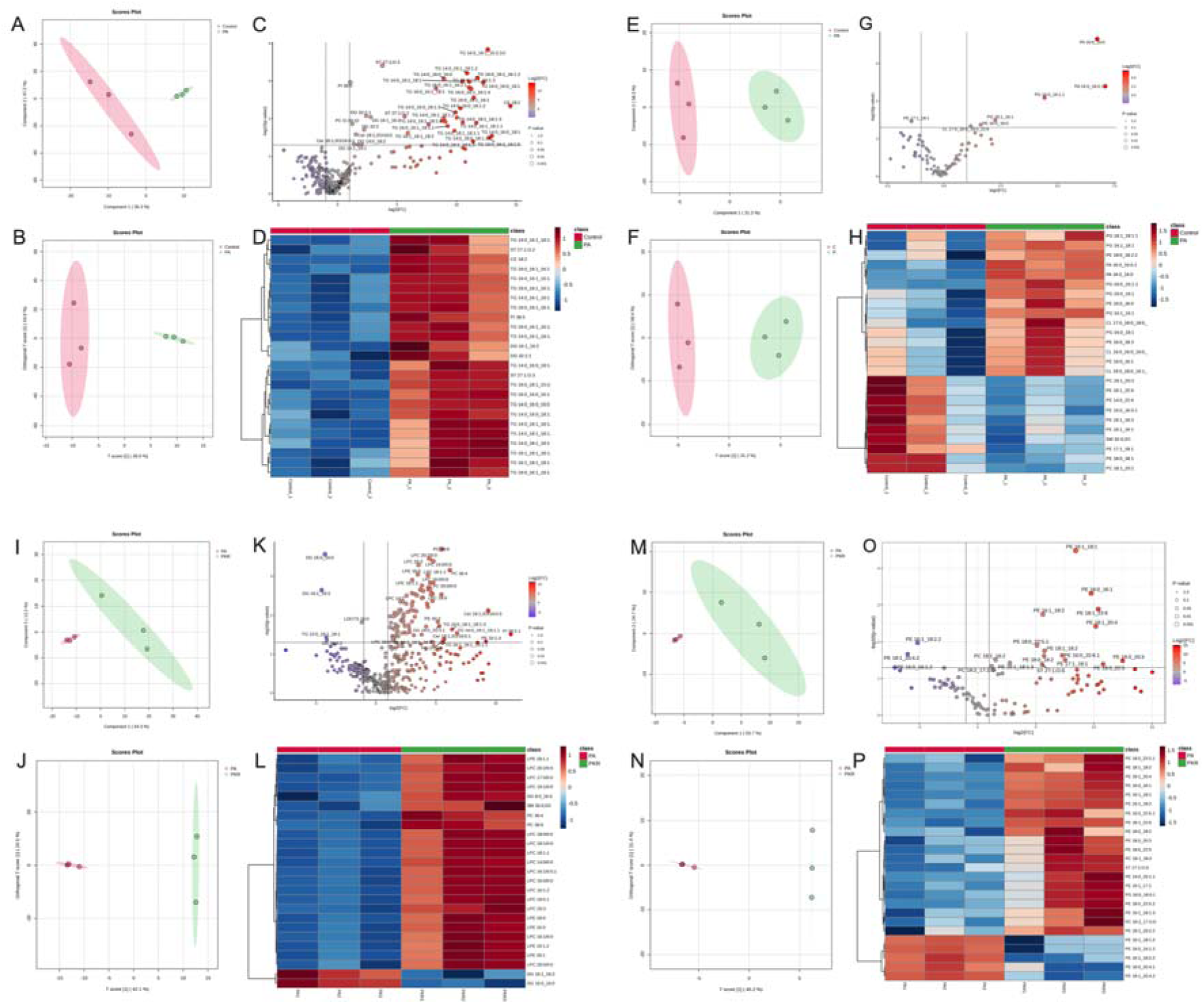
PKR Modulates Palmitate-Induced Lipidomic Alterations. PLS-DA score plots of Control and PA groups in positive (A) and negative (E) ion modes; OPLS-DA plots of control and PA in positive (B) and negative (F) modes. (C,G) Volcano plots and (D,H) heat maps; PLS-DA plots of PA and PKR groups in positive (I) and negative (M) ion modes; OPLS-DA plots for PA vs PKR comparison in positive (J) and negative (N) modes; (K,O) Volcano plots and (L,P) heat maps.

**Figure 4.**
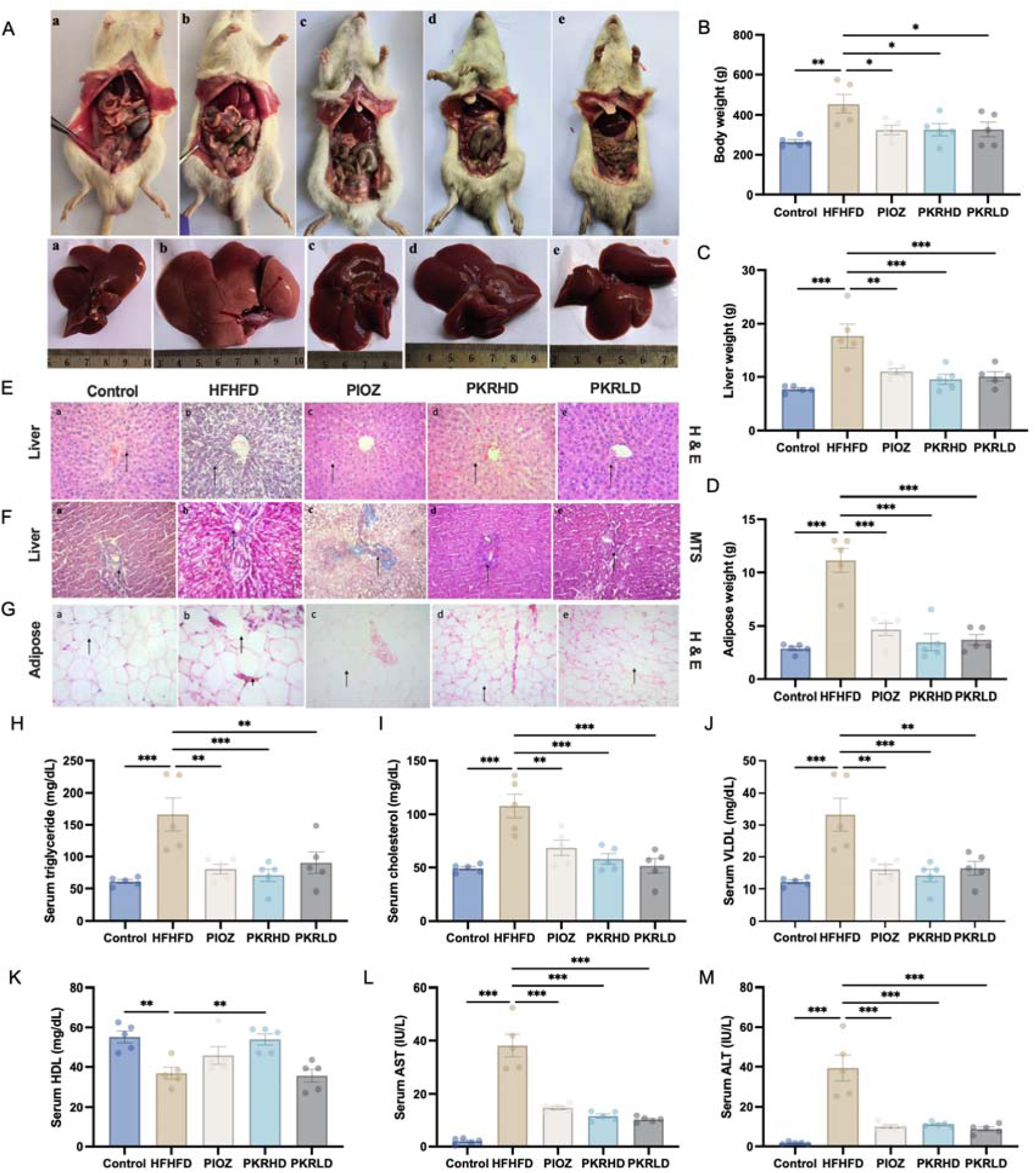
PKR mitigates HFD-induced hepatic steatosis, and dyslipidemia in rats. (A-B) Images of dissected rat with respective livers (a) control, (b) HFHFD, (c) PIOZ, (d) PKRHD and (e) PKRLD; (B)Body weight changes; (C) Liver weight; (D) Adipose tissue weight; (E & G) Representative H&E-stained sections of liver and adipose (400x); (F) Representative MTS stained sections of liver (400x); (H-K) Serum lipid parameters; (L) Serum AST levels; (M) Serum ALT levels. Data are presented as mean□±□SEM (n□=□5); *p□<□0.05, **p□<□0.01, ***p□<□0.001.

### 3.5. PKR enhances glucose homeostasis and incretin signalling with modest effects on pancreatic **β**-cell architecture

Histopathological evaluation of pancreatic tissue revealed a moderate improvement in islet number and β-cell mass in PKR-treated groups, while the overall islet architecture remained largely comparable to the HFHFD control (Fig. 5A). These observations suggest that PKR exerts limited direct effects on pancreatic structural restoration under the present conditions. Despite modest histological changes, PKR treatment significantly improved systemic glucose handling. Oral glucose tolerance testing (OGTT) demonstrated enhanced glucose clearance in treated animals, as reflected by a marked reduction in the area under the curve (AUC) compared to the HFHFD group (Fig. 5B–C), indicating improved glycaemic control. Mechanistically, this improvement was accompanied by a significant elevation in circulating glucagon-like peptide-1 (GLP-1) levels in PKR-treated animals (Fig. 5E), suggesting activation of incretin-mediated pathways. The enhancement of GLP-1 signalling highlights a potential gut–metabolic axis through which PKR contributes to improved glucose homeostasis. In contrast, serum insulin levels showed only a modest increase that did not reach statistical significance (Fig. 5D), indicating that the observed metabolic improvements are likely mediated through enhanced insulin sensitivity and incretin action rather than increased insulin secretion. Collectively, these findings demonstrate that PKR improves glucose homeostasis primarily through incretin signalling and metabolic regulation, with minimal direct effects on pancreatic structural remodelling.

**Figure 5.**
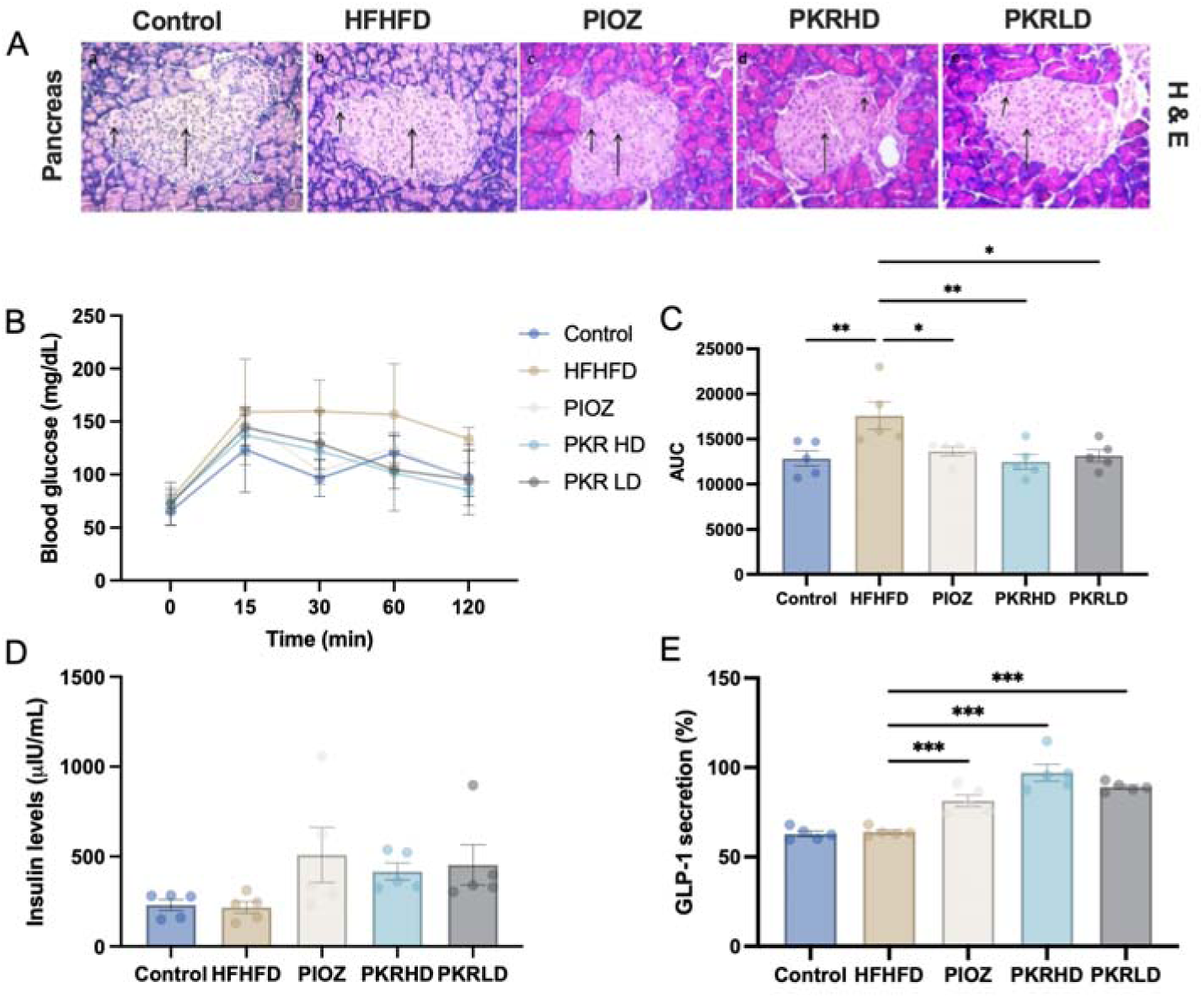
PKR improves glucose homeostasis and pancreatic function in HFD-induced MASLD via enhanced insulin and GLP-1 secretion. (A) Representative H&E-stained sections of pancreas (400x); (B) Oral Glucose Tolerance Test (OGTT) showing blood glucose levels at 0, 15-, 30-, 60-, and 120-minutes post-glucose administration; (C) Area Under the Curve (AUC) analysis of OGTT; (D) Plasma insulin levels; (E) Plasma GLP-1 levels. All data are presented as mean□±□SEM (n□=□5); *p□<□0.05, **p□<□0.01, ***p□<□0.001.

## 4. Discussion

Metabolic dysfunction is a complex, multifactorial condition characterized by dysregulated lipid metabolism, chronic low-grade inflammation, insulin resistance, and impaired energy homeostasis. Increasing evidence highlights lipotoxicity and membrane lipid imbalance as central drivers of metabolic derangements linking obesity, hepatic steatosis, and systemic metabolic impairment (18, 19). In this context, multi-target therapeutic strategies capable of modulating interconnected metabolic pathways are critically needed. The present study demonstrates that Patolakaturohiniyadi Kashayam (PKR), a classical Ayurvedic formulation, exerts coordinated regulatory effects on lipid metabolism, inflammatory signalling, and metabolic homeostasis, thereby addressing key pathological features of metabolic dysfunction. One of the central mechanistic insights from this study is the modulation of lipotoxicity, particularly through the reduction of diacylglycerol (DG) accumulation. DGs are well-recognized mediators of insulin resistance, primarily through activation of protein kinase C (PKC) isoforms, which impair insulin receptor signalling and downstream metabolic pathways (20). Elevated DG levels have been strongly associated with hepatic insulin resistance and metabolic dysfunction (21). By attenuating DG accumulation, PKR may alleviate lipotoxic stress and restore insulin signalling pathways, thereby improving metabolic outcomes. This aligns with previous findings demonstrating that targeting lipid intermediates, rather than total lipid content alone, is critical for reversing metabolic dysfunction (22).

Another important aspect of PKR action is its ability to restore phospholipid homeostasis, particularly the balance between phosphatidylcholine (PC) and phosphatidylethanolamine (PE). The PC/PE ratio is a key determinant of membrane integrity, mitochondrial function, and cellular homeostasis (23). Disruption of this balance has been implicated in metabolic diseases, including steatosis and insulin resistance (24). Enhanced PE levels, as observed in this study, are associated with improved mitochondrial dynamics and membrane curvature, which are essential for efficient lipid metabolism and energy production (25). Furthermore, increased lysophospholipids may reflect enhanced phospholipid turnover and membrane remodelling, processes that are critical for maintaining cellular adaptability under metabolic stress (26). Together, these findings suggest that PKR promotes a hepatoprotective lipid environment by reprogramming membrane lipid composition.

Inflammation represents another critical axis in the progression of metabolic dysfunction. Chronic activation of inflammatory pathways, particularly those involving TNF-α, IL-6, and NF-κB signalling, contributes to insulin resistance, adipose tissue dysfunction, and metabolic imbalance (27). Network pharmacology analysis in this study identified key inflammatory mediators, including TNF, IL6, and STAT3, as central targets of PKR. Modulation of these pathways may underlie the observed improvements in metabolic parameters. In particular, inhibition of TNF-α signalling has been shown to improve insulin sensitivity and reduce lipid accumulation (28), while suppression of IL-6/STAT3 signalling attenuates inflammatory responses and metabolic stress (29). The multi-component nature of PKR likely enables simultaneous targeting of these interconnected inflammatory pathways, thereby providing a broader therapeutic effect compared to single-target agents. Adipose tissue plays a pivotal role in systemic metabolic regulation, acting as both an energy storage organ and an endocrine tissue. Dysfunctional adipose tissue is characterized by impaired lipid buffering, increased lipolysis, and secretion of pro-inflammatory cytokines, which collectively contribute to ectopic lipid deposition and metabolic dysregulation (30). Therapeutic strategies that preserve adipose tissue function and prevent inflammatory remodelling are therefore essential for maintaining metabolic homeostasis. The ability of PKR to maintain adipose tissue integrity suggests a protective effect against adipose dysfunction, which may limit the release of free fatty acids and reduce lipid overflow to peripheral organs. This mechanism is particularly relevant in preventing the propagation of metabolic dysfunction across tissues. In addition to lipid and inflammatory pathways, incretin signalling has emerged as a critical regulator of metabolic homeostasis. Glucagon-like peptide-1 (GLP-1) plays a key role in glucose regulation by enhancing insulin secretion, suppressing glucagon release, and improving insulin sensitivity (31). Beyond its glycaemic effects, GLP-1 has been shown to influence lipid metabolism, reduce hepatic steatosis, and exert anti-inflammatory actions (32). The observed elevation of GLP-1 levels following PKR treatment suggests activation of the gut–metabolic axis, which may contribute to improved glucose and lipid homeostasis. Notably, the absence of a significant increase in insulin levels indicates that PKR may enhance metabolic efficiency primarily through incretin-mediated pathways and improved insulin sensitivity rather than direct stimulation of insulin secretion. This distinction is important, as therapies that improve insulin sensitivity without inducing hyperinsulinemia are generally associated with better long-term outcomes (33). The integration of network pharmacology with experimental validation provides further insight into the multi-target nature of PKR. Key hub genes identified in this study, including TP53, AKT1, and MAPK3, are involved in the regulation of cell survival, metabolism, and stress responses. The AKT signalling pathway, in particular, plays a central role in glucose uptake, lipid synthesis, and energy metabolism (34), while MAPK signalling is involved in inflammatory and stress responses (35). Modulation of these pathways suggests that PKR exerts a systems-level regulatory effect, targeting multiple nodes within the metabolic network. Such an approach is particularly advantageous in complex diseases like metabolic dysfunction, where single-target interventions often fail to achieve sustained therapeutic efficacy.

Traditional herbal formulations such as PKR offer a unique advantage in this regard due to their inherent polypharmacology. The presence of multiple bioactive compounds enables simultaneous modulation of diverse molecular targets, resulting in synergistic therapeutic effects (36). Several phytoconstituents present in PKR ingredients have been independently reported to exhibit lipid-lowering, anti-inflammatory, and hepatoprotective properties, supporting the observed pharmacological effects (37). However, the present study extends these findings by providing a comprehensive mechanistic framework linking these effects to lipidomic remodelling and metabolic regulation.

Despite these promising findings, certain limitations should be acknowledged. While the study provides strong evidence of metabolic regulation, further investigations are required to delineate the precise molecular interactions between PKR constituents and specific targets. Additionally, clinical studies are necessary to validate these findings in human populations and to establish the translational potential of PKR.

In conclusion, the present study demonstrates that PKR exerts multi-target therapeutic effects on metabolic dysfunction through coordinated regulation of lipid metabolism, inflammation, and incretin signalling. By attenuating lipotoxic intermediates, restoring phospholipid balance, and enhancing metabolic signalling pathways, PKR promotes systemic metabolic homeostasis. These findings highlight its potential as a promising multi-target intervention for the management of metabolic disorders, particularly those driven by lipid dysregulation.

## Author Contributions

Conceptualization: CNVP, SK and SKK. Methodology: SK. Validation: SK, CNVP and SKK. Formal analysis: SK. Investigation: SK. Resources: CNVP, PD and SKK. Data curation: SK. Writing – Original Draft: SK. Writing – Review & Editing: CNVP, PD and SKK. Visualization: CNVP, SKK and SK. Supervision: CNVP and SKK. Project administration: CNVP and SKK. Funding acquisition: CNVP and SKK.

## Ethical statement

The study protocol was approved by the institutional animal ethics committee (IAEC/ABMRCP/2023-2024/4).

## Acknowledgment

The authors also thank the University of Trans-Disciplinary Health Sciences and technology (TDU), Bangalore, for providing the necessary infrastructure and facilities to carry out the experiments and the Rural India Support Trust (RIST) for financial support. The authors also acknowledge Dr. Manjunatha P. Mudagal and Dr. Suresh Janadri for their guidance and support during the preclinical studies. Figures generated with Biorender.com.

## Data availability

The data associated with this article are provided within this article and as a supplementary material. The data can also be requested by the corresponding author.

## Declaration of generative AI in scientific writing

The authors declare that this article did not use generative AI or AI-assisted technologies.

## References

1. World Health Organization. Obesity and overweight. 2023.

2. Ahirwar R, Mondal PR. Prevalence of obesity in India: A systematic review. Diabetes Metab Syndr. 2019;13(1):318–321.

3. Pradeepa R, Anjana RM, Joshi SR, et al. Prevalence of generalized & abdominal obesity in urban & rural India. Indian J Med Res. 2015;142(2):139–150.

4. Younossi ZM, Koenig AB, Abdelatif D, et al. Global epidemiology of NAFLD. Hepatology. 2016;64(1):73–84.

5. Duseja A, et al. Non-alcoholic fatty liver disease in India – a lot done, yet more required. Indian J Gastroenterol. 2010;29(6):217–225.

6. Powell EE, Wong VW, Rinella M. NAFLD: metabolic implications and complications. Lancet. 2021;397(10290):2212–2224.

7. Sun K, Tordjman J, Clément K, Scherer PE. Fibrosis and adipose tissue dysfunction. Cell Metab. 2013;18(4):470–477.

8. Buzzetti E, Pinzani M, Tsochatzis EA. NAFLD pathogenesis. Metabolism. 2016;65(8):1038–1048.

9. Samuel VT, Shulman GI. Mechanisms of insulin resistance. Cell. 2012;148(5):852–871.

10. Romero-Gómez M, Zelber-Sagi S, Trenell M. Treatment of NAFLD with diet and exercise. J Hepatol. 2017;67(4):829–846.

11. Friedman SL, Neuschwander-Tetri BA, Rinella M, Sanyal AJ. Mechanisms of NAFLD and therapeutic targets. Nat Med. 2018;24(7):908–922.

12. Sharma PV. Dravyaguna Vijnana. Chaukhambha Bharati Academy; 2001.

13. Ainsworth EA, Gillespie KM. Estimation of total phenolic content and other oxidation substrates in plant tissues using Folin-Ciocalteu reagent. Nat Protoc. 2007;2(4):875–877. doi:10.1038/nprot.2007.102

14. Thottappillil A, Sahoo S, Chakraborty A, et al. In vitro and in silico analysis proving DPP4 inhibition and diabetes-associated gene network modulation by a polyherbal formulation: Nisakathakadi Kashaya. J Biomol Struct Dyn. 2024;42(24):13588–13602. doi:10.1080/07391102.2023.2276880

15. Bligh EG, Dyer WJ. A rapid method of total lipid extraction and purification. Can J Biochem Physiol. 1959;37(8):911–917. doi:10.1139/O59-099

16. OECD. Test No. 425: Acute Oral Toxicity: Up-and-Down Procedure. OECD Guidelines for the Testing of Chemicals, Section 4. Published online June 29, 2022. doi:10.1787/9789264071049-EN

17. Ghosh M. N. Fundamentals of Experimental Pharmacology, Hilton & Company, 7^th^ Edition, 2020.

18. Samuel VT, Shulman GI. Mechanisms for insulin resistance: common threads and missing links. Cell. 2012;148(5):852–871.

19. Buzzetti E, Pinzani M, Tsochatzis EA. The multiple-hit pathogenesis of non-alcoholic fatty liver disease (NAFLD). Metabolism. 2016;65(8):1038–1048.

20. Itani SI, Ruderman NB, Schmieder F, Boden G. Lipid-induced insulin resistance in human muscle is associated with changes in diacylglycerol, protein kinase C, and I_κB_-_α_. J Clin Invest. 2002;110(2):181–190.

21. Petersen MC, Shulman GI. Mechanisms of insulin action and insulin resistance. Cell Metab. 2018;27(1):22–35.

22. Farese RV Jr, Zechner R, Newgard CB, Walther TC. The problem of establishing relationships between hepatic steatosis and hepatic insulin resistance. Science. 2012;337(6090):1496–1499.

23. van der Veen JN, Kennelly JP, Wan S, Vance JE, Vance DE, Jacobs RL. The critical role of phosphatidylcholine and phosphatidylethanolamine metabolism in health and disease. Biochim Biophys Acta Biomembr. 2017;1859(9 Pt B):1558–1572.

24. Li Z, Vance DE. Phosphatidylcholine and choline homeostasis. J Biol Chem. 2008;283(4):2225–2229.

25. Tasseva G, Bai HD, Davidescu M, Haromy A, Michelakis E, Vance JE. Phosphatidylethanolamine deficiency in mammalian mitochondria impairs oxidative phosphorylation and alters mitochondrial morphology. J Biol Chem. 2013;288(6):415–426.

26. Law SH, Chan ML, Marathe GK, Parveen F, Chen CH, Ke LY. An updated review of lysophosphatidylcholine metabolism in human diseases. Int J Mol Sci. 2019;20(5):1462.

27. Hotamisligil GS. Inflammation and metabolic disorders. Nature. 2006;444(7121):860–867.

28. Kern PA, Ranganathan S, Li C, Wood L, Ranganathan G. Adipose tissue tumor necrosis factor and interleukin-6 expression in human obesity and insulin resistance. Diabetes. 2001;50(10):2306–2313.

29. Heinrich PC, Behrmann I, Haan S, Hermanns HM, Müller-Newen G, Schaper F. Principles of interleukin (IL)-6-type cytokine signalling and its regulation. Biochem J. 2003;374(Pt 1):1–20.

30. Sun K, Tordjman J, Clément K, Scherer PE. Fibrosis and adipose tissue dysfunction. Cell Metab. 2013;18(4):470–477.

31. Drucker DJ. The biology of incretin hormones. Cell Metab. 2006;3(3):153–165.

32. Nauck MA, Quast DR, Wefers J, Meier JJ. GLP-1 receptor agonists in the treatment of type 2 diabetes – state-of-the-art. Diabetes Care. 2021;44(1):234–243.

33. DeFronzo RA. From the triumvirate to the ominous octet: a new paradigm for the treatment of type 2 diabetes mellitus. Diabetes. 2009;58(4):773–795.

34. Manning BD, Toker A. AKT/PKB signaling: navigating the network. Cell. 2017;169(3):381–405.

35. Cargnello M, Roux PP. Activation and function of the MAPKs and their substrates. Microbiol Mol Biol Rev. 2011;75(1):50–83.

36. Hopkins AL. Network pharmacology: the next paradigm in drug discovery. Nat Chem Biol. 2008;4(11):682–690.

37. Gupta R, Sharma AK, Dobhal MP, Sharma MC, Gupta RS. Antidiabetic and antioxidant potential of herbal formulations: a review. J Ethnopharmacol. 2017;206:45–56.

