## Supplementary Table 1 for "*Patolakaturohiniyadi Kashayam*, exerts anti-steatotic and anti-obesogenic effects via coordinated regulation of lipid metabolism, inflammation, and incretin signalling"

Table 1. Composition of HFHFD and chow diet

| **Composition of the HFHFD Diet** | | |
| --- | --- | --- |
| **Sl. No** | **Particulars** | **Value** |
| 1 | Protein | 18.15% |
| 2 | Fat | 35.10% |
| 3 | Fiber | 2.5% |
| 4 | Carbohydrates | 20% |
| 5 | Calcium | 1.25% |
| 6 | Total ash | 3.5% |
| 7 | Phosphorous | 0.67% |
| 8 | Energy through fat | 3120 kcal/kg (60%) |
| 9 | Energy through protein | 1050 kcal/kg (20%) |
| 10 | Energy through carbohydrates | 1030 kcal/kg (20%) |
| 11 | Energy from palm oil | 2700 kcal/kg (52%) |
| 12 | Moisture | 2.5% |
| 13 | Cholesterol | 2% |
| 14 | Cholic acid | 0.5% |
| 13 | Energy | 5200 kcal/kg |
| **Composition of the Chow Diet** | | |
| **Sl. No** | **Particulars** | **Value** |
| 1 | Crude protein | 7.47% |
| 2 | Fat | 3.25% |
| 3 | Fiber | 2.81% |
| 4 | Carbohydrates | 63% |
| 5 | Calcium | 1.25% |
| 6 | Total ash | 4.7% |
| 7 | Phosphorous | 0.66% |
| 8 | Moisture | 7.47% |
| 9 | Energy | 3060 kcal/kg |
